# HIV-1 Env induces pexophagy and an oxidative stress leading to uninfected CD4+ T cell death

**DOI:** 10.1101/2019.12.19.881110

**Authors:** Coralie F Daussy, Mathilde Galais, Baptiste Pradel, Véronique Robert-Hebmann, Sophie Sagnier, Sophie Pattingre, Martine Biard-Piechaczyk, Lucile Espert

## Abstract

The immunodeficiency observed in HIV-1-infected patients is mainly due to uninfected bystander CD4+ T lymphocytes death. The viral envelope glycoproteins (Env), expressed at the surface of infected cells, play a key role in this process. Env triggers autophagy, process necessary to subsequent apoptosis, and to production of Reactive Oxygen Species (ROS) in bystander CD4+ T cells. Here, we demonstrate that Env-induced oxidative stress is responsible for their death by apoptosis. Moreover, we report that peroxisomes, organelles involved in the control of oxidative stress, are targeted by Env-mediated autophagy. Indeed, we observe a selective autophagy-dependent decrease in the expression of peroxisomal proteins, catalase and PEX14, upon Env exposure, since the down-regulation of either BECLIN 1 or p62/SQSTM1 restores their expression levels. Fluorescence studies allowed us to conclude that Env-mediated autophagy degrades these entire organelles and specifically the mature ones. Together, our results on Env-induced pexophagy provide new clues on HIV-1-induced immunodeficiency.

## Introduction

HIV-1 infection is characterized by a progressive decline in the number of CD4+ T lymphocytes, ultimately leading to AIDS (Acquired Immuno-Deficiency Syndrome), in untreated patients^1, 2^. Importantly, the majority of dying CD4+ T cells are not productively infected but are bystander uninfected cells.^3–6^. Among the several mechanisms identified to explain this depletion^7–10^, it has been demonstrated that HIV-1 envelope glycoproteins, composed of gp120 and gp41 associated in trimers (here called Env), play a major role. Indeed, Env expressed at the surface of infected cells is able to induce apoptosis in bystander CD4+ T cells after its interaction with the CD4 receptor and the co-receptor, either CCR5 or CXCR4^11,12^. Our team has demonstrated that Env, and specifically the gp41 fusogenic function, triggers a massive induction of macroautophagy (hereafter referred to as autophagy) in bystander CD4+ T cells^13, 14^. Importantly, this mechanism is necessary for the subsequent apoptosis, suggesting that Env-mediated autophagy could specifically degrade a component crucial for cellular survival^14^. Interestingly, we and others, have also shown that Env triggers an oxidative stress in bystander CD4+ T cells, characterized by a strong production of Reactive Oxygen Species (ROS)^15, 16^.

Autophagy is a catabolic pathway involved in many cellular functions^17^. It can be divided in three major steps: (i) the initiation step, with the formation of a double membrane structure named phagophore, (ii) the elongation step, where this structure elongates engulfing portions of cytoplasm and then closes to form a vacuole called autophagosome, and (iii) the maturation step, where the autophagosomes fuse with lysosomes to degrade and recycle the sequestered material. More than 30 genes and several signalling platforms are involved in autophagosomes formation, such as the Class III PI3kinase/BECLIN 1 complex, acting at the initiation step of the process, and two ubiquitination-like conjugation systems leading to the covalent conjugation of ATG8 proteins to a lipid, the Phosphatidyl-Ethanolamine (PE), involved in the elongation step of autophagosomal membranes^18^. Autophagic degradation can be highly selective through proteins called autophagy receptors, such as p62/SQSTM1 (Sequestosome 1)^19^. These proteins bind to both the PE-conjugated ATG8 proteins and to substrates, allowing their targeting to autophagosomes^20^.

Autophagy and oxidative stress are intimately connected and the level of (ROS) production is tightly regulated. Low amounts of ROS are involved in the activation of survival signalling pathways^21^, whereas an excessive ROS production activates cell death^22^. By degrading organelles involved in ROS production, such as damaged mitochondria, selective autophagy counterbalances excessive ROS production^23^. However, when the levels of ROS become excessive, cell death is initiated either by apoptosis or necrosis^22^. Alternatively, autophagy controls the cellular oxidative status through its involvement in peroxisomes homeostasis. Peroxisomes are essential single membrane-limited organelles containing multiple enzymes involved in metabolic and antioxidant reactions. Catalase, one of the major peroxisomal enzyme, catalyses the transformation of hydrogen peroxide (H_2_O_2_) into water, therefore detoxifying the cell from this harmful component^24, 25^. Peroxins (PEX proteins) and PMP (Peroxisomal Membrane Proteins) control peroxisome assembly and division. Peroxisomes acquire their various components along their biogenesis and grow towards mature peroxisomes, reaching a size between 0.5μm and 1μm. Selective autophagy of peroxisomes, called pexophagy, is responsible for the number and quality of these organelles^26^ by maintaining an equilibrium between their biogenesis and breakdown^27, 28^.

The aim of the present study was to understand how Env-mediated autophagy could trigger apoptosis in bystander CD4+ T cells. Given that Env also induced an accumulation of ROS in target cells and considering the essential role of peroxisomes in the regulation of the oxidative status of the cell, we focused our research on the interplay between autophagy and peroxisomes upon Env exposure. Our results show that Env-mediated autophagy is responsible for ROS accumulation in bystander CD4+ T cells and demonstrate that Env-mediated autophagy selectively targets mature peroxisomes.

## Results

### Env-mediated autophagy and ROS production are responsible for bystander CD4+ T cells apoptosis

We previously demonstrated that Env induces autophagy in target bystander CD4+ T cells, ultimately leading to apoptosis^29^. These results were obtained with different types of effector cells and target cells also used in the present study (Figure 1A). These models, based on the co-culture between effector cells expressing Env and target cells that express CD4 and CXCR4, allow the analysis of the Env effect in absence of viral replication. In the model 1, the target cells were either primary CD4+ T lymphocytes or the CEM-derived A201/CD4.403 T cell line. These target cells were co-cultured with HEK293 stably expressing Env (HEK.Env) during 48h. The parental HEK293 cell line was used as a negative control. In the model 2, the target cells were HEK293 cells stably expressing CD4.403 and CXCR4 and are referred to as (HEK/CD4.403/CXCR4). These target cells were co-cultured for 48h with 8.E5 cells expressing Env and harbouring a single defective copy of replication incompetent HIV-1 proviral DNA. The parental CEM cell line was used as a negative control. Importantly, Env-induced apoptosis and its dependence on autophagy are equivalent in these two co-culture models.

**Figure 1:**
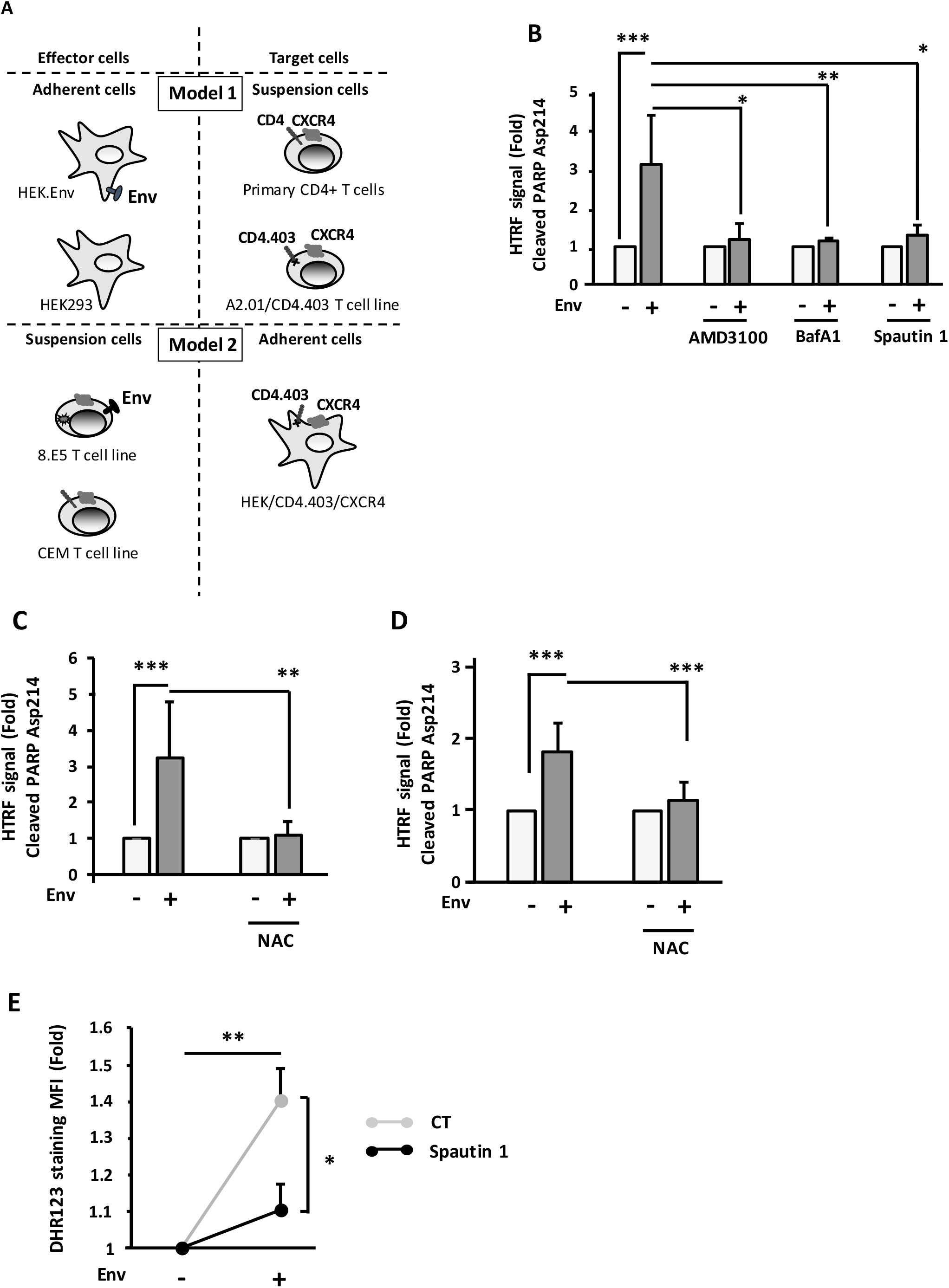
Env-mediated autophagy participated in ROS production that is responsible for bystander CD4+ T cell apoptosis. **A.** Description of the cellular models. The experiments are based on the co-culture between effector cells expressing Env at their surface and target cells expressing CD4 and CXCR4. In the model 1, target cells are either primary CD4+ T lymphocytes purified from healthy donor blood samples or a CD4+ T cell line expressing CXCR4. Effector cells are HEK293 stably expressing Env (HEK.Env). HEK293 cells are used as negative control. In the model 2, target cells are HEK.293 cells stably expressing CD4 and CXCR4. Effector cells are 8.E5 cells, which are chronically infected cells and thus express Env at their surface. The parental CEM cell line is used as a negative control. **B.** Env-induced apoptosis depends on autophagy. Target CD4+ T cells were co-cultured for 48h with HEK293 or HEK.Env in the presence or absence of AMD3100 (1μg/ml), BafA1 (50 nM) or Spautin 1 (50 μM). Target cells were harvested and fold increase in Env-mediated apoptosis was studied by quantifying the cleavage of PARP using HTRF. Results are from at least 3 independent experiments. *p<0.05; **p < 0.01; ***p < 0.001. **C. and D.** Target CD4+ T cells (either T cell line (C), or primary CD4+ T cells (D)) were co-cultured for 48h with HEK293 or HEK.Env in the presence or absence of NAC (10 mM). Target cells were harvested and fold increase in Env-mediated apoptosis was studied by quantifying the cleavage of PARP using HTRF. Data are representative of at least three independent experiments. **p < 0.01; ***p < 0.001 **E.** Target CD4+ T cells were co-cultured for 48h with HEK.293 or HEK.Env in the presence or absence of Spautin 1 (50 μM). Analysis of intracellular ROS production was measured by flow cytometry using DHR 123. Data are representative of at least three independent experiments. *p<0.05; **p < 0.01.

Using the model 1, we confirm our previous results using an HTRF method (Homogeneous Time Resolved Fluorescence), allowing the quantitative detection of the cleaved PARP (poly (ADP-ribose) polymerase), which is a known caspase-3 substrate during apoptosis. Env-induced cleavage of PARP is completely blocked in the presence of AMD3100, which is an antagonist of CXCR4^30^, demonstrating that apoptosis is due to Env binding to this co-receptor (Figure 1B). Moreover, Env-mediated apoptosis is dependent on autophagy since it is abolished by addition of Spautin 1 and Bafilomycin A1 (BafA1), which are inhibitors of the early and late steps of autophagy respectively (Figure 1B).

We, and others, previously demonstrated that cell expressing Env induced an oxidative stress in bystander CD4+ T cells, characterized by a strong production of ROS^15, 16^. We thus evaluated the involvement of the Env-induced production of ROS in apoptosis. We co-cultured effector cells, expressing Env, with target CD4+ T cells (Model 1, Figure 2A), either a CD4+ T cell line (Figure 1C) or blood-purified primary CD4+ T cells (Figure 1D), in the presence of N-Acetyl-Cysteine (NAC), which is a general antioxidant molecule. Env-mediated accumulation of ROS is responsible for apoptosis since the addition of NAC completely abolished cell death.

**Figure 2:**
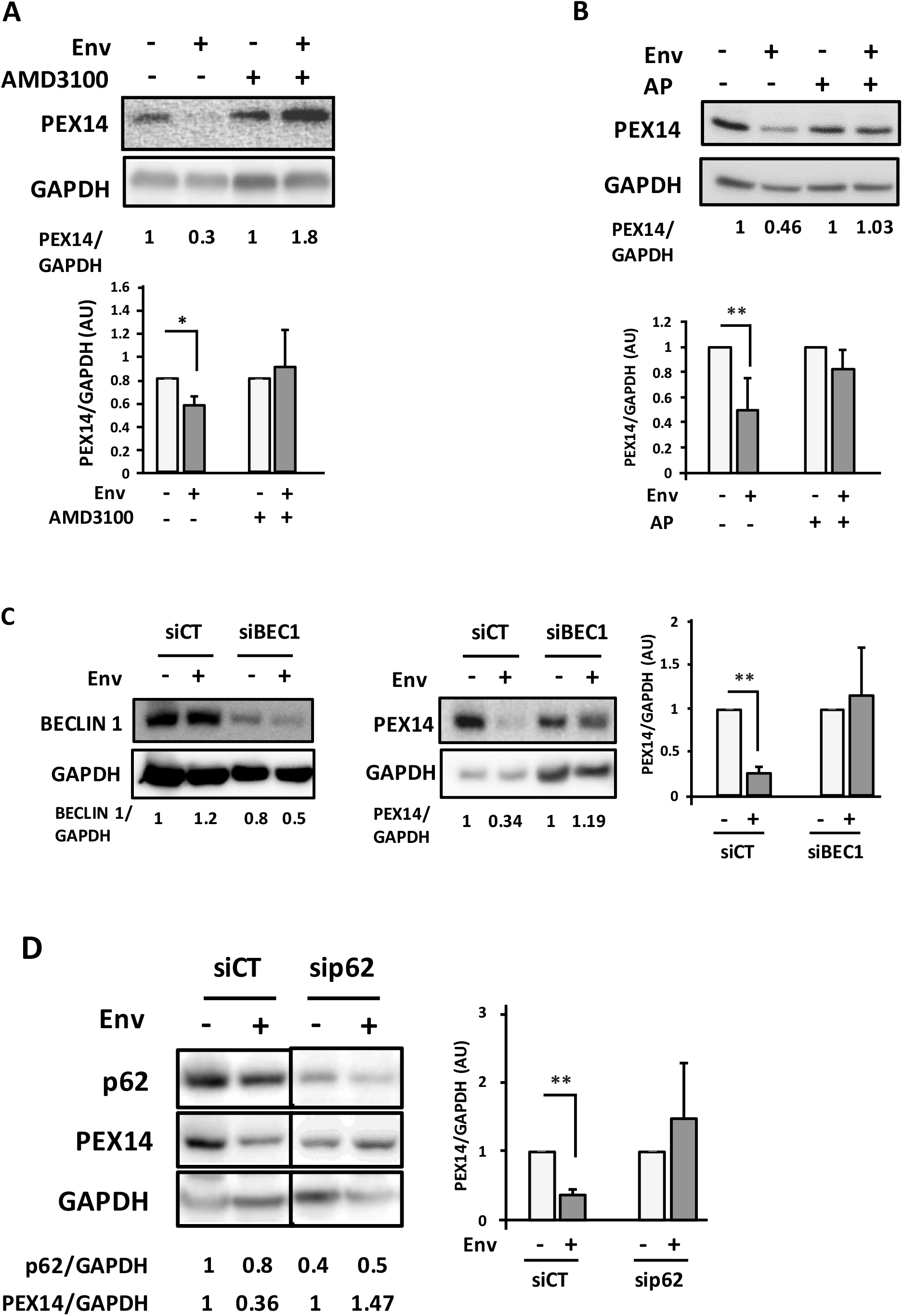
PEX14 is degraded by Env-induced autophagy. **A.** Env induces a decrease in PEX14 expression level. PEX14 expression level was analyzed in target CD4+ T cells after 48h of co-culture with effector HEK293 cells expressing or not Env, in the presence or absence of AMD3100 (1μg/ml). The expression level ratios of PEX14 were calculated between the conditions with or without Env and normalized to that obtained with anti-GAPDH Ab. Data are representative of at least 3 independent experiments. *p < 0.05. **B.** PEX14 is degraded in lysosomes in response to Env. PEX14 expression level was analyzed in target CD4+ T cells after 48h of co-culture with effector HEK293 cells expressing or not Env, in the presence or absence of AP (E64d + Pepstatin A, 10 μM each). The expression level ratios of PEX14 were calculated between the conditions with or without Env and normalized to that obtained with anti-GAPDH antibody. Data are representative of at least 3 independent experiments. **p < 0.01. **C.** Autophagy is involved in Env-mediated PEX14 degradation. HEK/CD4.403/CXCR4 cells were co-cultured during 48h with effector cells expressing or not Env (8.E5 or CEM, respectively) after their transfection with *Beclin 1* (siBEC1) or control (siCT) siRNAs. Reduction in BECLIN 1 expression was analyzed by western blot using the specific Abs. The expression level ratios of PEX14 were calculated between the conditions with or without Env and normalized to that obtained with anti-GAPDH Ab. Data are representative of at least 3 independent experiments. **p < 0.01. **D.** p62/SQSTM1 is involved in the Env-mediated PEX14 degradation. HEK/CD4.403/CXCR4 cells were co-cultured during 48h with effector cells expressing or not Env (8.E5 or CEM, respectively) after their transfection with *p62/SQSTM1* (sip62) or control (siCT) siRNAs. Reduction in p62/SQSTM1 expression was analyzed by western blot using the specific Abs. The expression level ratios of PEX14 were calculated between the conditions with or without Env and normalized to that obtained with anti-GAPDH Ab. Data are representative of at least 3 independent experiments. **p < 0.01.

The antioxidant properties of autophagy are well characterized. We thus analyzed the effect of an inhibition of this process on the Env-induced ROS production by using Spautin 1. After 48h of coculture (Model 1), we added DHR123 to the cells to quantify ROS. Surprisingly, we found that inhibition of autophagy leads to a decrease in ROS production (Figure 2B) strongly suggesting that autophagy is involved in the Env-mediated accumulation of ROS, which is responsible for bystander CD4+ T cell apoptosis.

### Autophagy selectively targets peroxisomal proteins

Peroxisomes are major cellular antioxidant organelles. Their number and integrity are regulated through selective degradation by autophagy in a process called pexophagy. We hypothesized that Env-induced autophagy could degrade peroxisomes in target cells, leading to the accumulation of ROS. To test this hypothesis, we analyzed the level of two peroxisomal proteins by western blot, in target cells after their co-culture during 48h with Env-expressing cells in different conditions. We followed the expression level of catalase, which is one of the major peroxisomal enzymes. As catalase is also present in mitochondria, we also analyzed PEX14, which is strictly located at the surface of peroxisomes^31^. As shown in Figure 2A and Supplementary Figure S1, the levels of PEX14 and catalase respectively decreased in target cells in contact with Env. AMD3100 rescued their level, confirming the involvement of Env in this effect. Next, to analyze the role of autophagy in Env-mediated PEX14 and catalase decreases, we added lysosomal protease inhibitors (AP for AntiProteases: E64d and Pepstatin A) to the co-culture. We observed that the Env-mediated PEX14 and catalase level decreases are mediated by lysosomal degradation (Figure 2B and Supplemental Figure S2). Notably, we verified that the observed diminution of PEX14 expression levels was not due to a difference in Pex14 mRNA production (Supplementary Figure S3).

To definitively attest the role of Env-induced autophagy on PEX14 and catalase levels, we inhibited autophagy by transfecting siRNA targeting the expression of BECLIN 1 in target cells before their co-culture with Env-expressing cells. We still observed a PEX14 decrease in target cells transfected with a control siRNA upon Env exposure, but PEX14 expression level remains unchanged when target cells, in which BECLIN 1 was downregulated, are co-cultured with Env-expressing cells (Figure 2C). Of note, similar results were obtained for catalase (Supplementary Figure S4). Taken together, these results confirmed that Env induces the autophagic degradation of peroxisomal proteins.

To complete the analysis, we checked the involvement of the p62/SQSTM1 autophagy receptor, known for its involvement in pexophagy. We transfected siRNA targeting p62/SQSTM1 in target cells and co-cultured them with Env-expressing cells. The PEX14 protein is no longer degraded in target cells where the expression of p62/SQSTM1 has been invalidated (Figure 2D). Altogether, these results show that Env-induced autophagy selectively target peroxisomal proteins in bystander CD4+ T cells.

### Env-induced autophagy targets mature peroxisomes in target cells

Based on the above results, we next wanted to analyze whether the entire peroxisomes are degraded by autophagy. We labelled peroxisomes in target cells with green fluorescent protein (GFP) by transfecting the GFP-SKL plasmid that contains a C-terminal tripeptide, SKL, called “Peroxisomal Targeting Signal” (PTS), targeting the GFP into peroxisomes. We confirmed that this construct stains peroxisomes by labelling PEX14 in the GFP-SKL-transfected cells. The PEX14 puncta massively co-localized with the GFP signal, confirming that the GFP-SKL construct efficiently labels peroxisomes (Figure 3A). We thus co-cultured the GFP-SKL-transfected target cells with Env-expressing cells during 24h and 48h and quantified the GFP signal using flow cytometry. We observed a significant decrease in the GFP signal in target cells in contact with Env-expressing cells in a time-dependent manner (Figure 3B). Interestingly, the fluorescence signal is rescued by addition of Spautin 1, arguing that Env induces pexophagy. To further confirm this result, we analyzed the co-localisation between GFP-SKL and MAP-LC3B (here called LC3B), which is an ortholog of ATG8, by immunofluorescence. Target cells, transfected with the GFP-SKL construct, were co-cultured during 48h with Env-expressing cells in presence of BafA1 to block the autophagic flux. By confocal microscopy, we observed several co-localisations between LC3B and GFP-SKL (Figure 3C). As the number of peroxisomes and autophagosomes was very high in the conditions used, we decided to quantify the co-localisation events between GFP-SKL and LC3B in the total target cell volume. To this aim, several optical slices of target cells (Z=1μm) were acquired after their co-culture with effector cells expressing or not Env. We used the ImageJ Plugin “JaCOP” allowing the determination of the Mander’s co-localisation score between LC3B puncta and peroxisomes. We observed a significant increase of GFP-SKL colocalizing with LC3B in target cells after their contact with Env-expressing cells compared to the control (Figure 3C), confirming that peroxisomes are targeted to autophagy in cells exposed to Env.

**Figure 3:**
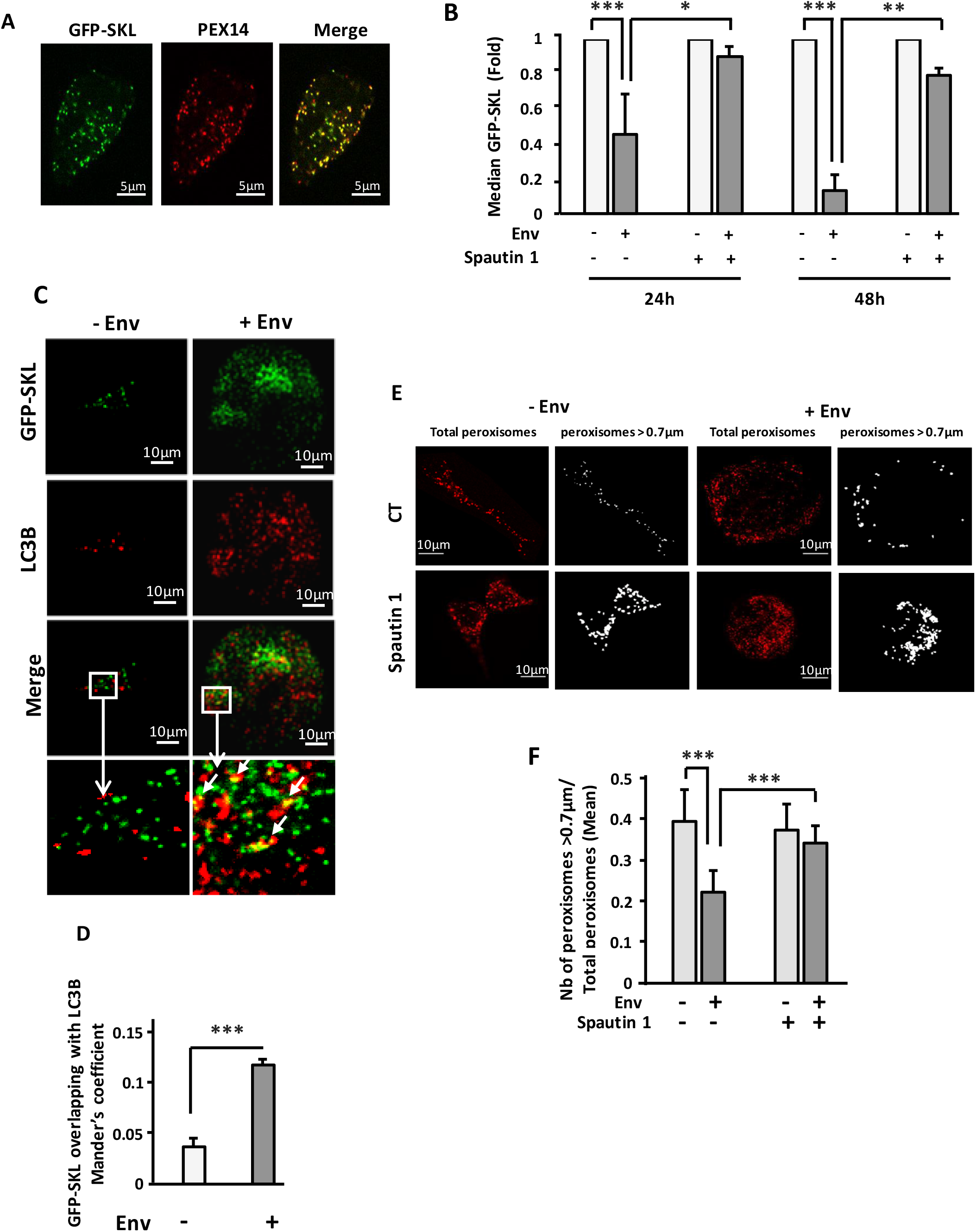
Env-induced autophagy targets mature peroxisomes. **A.** The SKL motif drives GFP to peroxisomes. Target cells were transfected with the GFP-SKL construct. 24h later, peroxisomes were labelled using an anti-PEX14 antibody. Images were acquired with a confocal microscope. **B.** The GFP labelled peroxisomes decrease overtime in target cells after their co-culture with effector cells expressing Env. Target cells were transfected with the GFP-SKL construct. 24h later, the transfected cells were co-cultured with effector cells in presence or not of Spautin 1 (50 μM). The Mean Fluorescence Intensity of the GFP signal was analyzed in target cells using flow cytometry after 24 and 48h of co-culture. Data are representative of at least 3 independent experiments. *p<0.05; **p < 0.01; ***p < 0.001. **C.** The colocalization between the GFP labelled peroxisomes and LC3B increases in target cells upon Env exposure. Target cells were transfected with the GFP-SKL construct. 24h later, the transfected cells were co-cultured with effector cells in the presence of BafA1 (50nM). Autophagosomes were labelled using an anti-LC3B by immunofluorescence. **D.** The graph represents the overlap of peroxisomes staining in the LC3B-positive structures. Data are representative of at least 3 independent experiments. ***p < 0.001. **E.** Target cells were stained by immunofluorescence with an anti-PEX14 antibody after 48h of co-cultures with effector cells in presence or not of Spautin 1 (50 μM). The peroxisomes were sorted according to their size using the “Cell Profiler software”. **F.** The total volume of the target cells was imaged using a confocal microscope. Data obtained with the “Cell Profiler software” are representative of at least 3 independent experiments. ***p < 0.001.

As peroxisomes respond to changes in the cellular environment by adapting their number and morphology^32^, we analyzed these last parameters in target cells after their contact with Env-expressing cells. We used DAB (3’3-DiAminoBenzidine) to label peroxisomes in targets cells after 48h of co-culture with effector cells in the presence or absence of AP to block the autophagic flux. DAB allows the observation of peroxidase activity by inducing a black precipitate in the cells^33^. Surprisingly, we observed, by light microscopy, different aspects of DAB precipitates depending on the presence of Env. Indeed, the precipitates seemed smaller in target cells co-cultured with Env-expressing cells. The addition of lysosomal protease inhibitors led to the restoration of bigger precipitates. These observations suggested that only the mature peroxisomes were degraded through autophagy (Supplementary Figure S5). As the images resolution did not allow us to quantify the DAB precipitates of each size in the different conditions, we decided to confirm and quantify these results by analyzing peroxisomes in target cells by immunofluorescence with a particular focus on their size. To this aim, we stained peroxisomes in target cells after 48h of co-culture with Env-expressing cells using an anti-PEX14 antibody in presence or absence of Spautin 1. Peroxisomes vary in size from 0.1 to 1 μm in diameter. Although they are capable of fission, the “mature” peroxisomes, which exert their antioxidant capacity by catalase, are larger than those undergoing biogenesis. The target cells were analyzed by confocal microscopy (Figure 3E). In order to distinguish large peroxisomes from small ones (in the process of biogenesis), we used the “Cell Profiler” software and designed a protocol allowing us to measure their area in target cells. We fixed a “Z” of 1μm between each optical slice to avoid acquiring twice the same peroxisome. Each acquisition, with a confocal microscope, was performed on the total volume of the target cells. We set up a threshold of 0.7 μm in diameter to identify “mature” peroxisomes. Based on these measurements, we established, for each image analyzed, a ratio between the number of mature peroxisomes (diameter> 0.7μm) and the number of total peroxisomes. The data were collected for each image constituting the volume of a target cell. Interestingly, this ratio is decreased in target cells after their contact with Env-expressing cells and the inhibition of autophagy, by addition of Spautin1, restores the ratio close to the one of control cells (Figure 3F). These results strongly suggest that Env-induced autophagy degrades only mature peroxisomes.

## Discussion

The interplay between autophagy and HIV-1 infection is complex and depends on the target cell type and on the infectious status of the cell (infected *vs* bystander cells)^34–36^. We previously showed that Env, expressed on infected cells, induces autophagy in bystander CD4+ T cells, after its interaction with HIV-1 receptors present at their surface. Notably, this Env-induced autophagy leads to apoptosis of these target cells, mechanism likely contributing to CD4+ T cells depletion observed in HIV-1 associated physiopathology^14^. As HIV-1 Env induces the production of ROS in bystander CD4+ T cells^16^, we first demonstrated that this oxidative stress is responsible for subsequent apoptosis. The cellular ROS levels are tightly regulated by the action of antioxidant enzymes in order to avoid their harmful effect. As we previously showed that Env-mediated autophagy is a prerequisite to apoptosis in bystander cells, we hypothesized that antioxidant pathways are targeted by autophagy and, consequently, deprive the cell of their protective action. Inhibition of autophagy by adding Spautin1 blocks apoptosis (Figure 1B), but also dramatically decreases the production of ROS in bystander cells cocultured with effector cells expressing Env (Figure 2C), strongly suggesting that Env-induced autophagy leads to an uncontrolled oxidative stress in bystander cells. ROS can be produced at different cellular sites and three organelles are known to control it: mitochondria, endoplasmic reticulum and peroxisomes, forming the “redox triangle”^37^. These three organelles are interconnected and cooperate to fight oxidative stress. Importantly, the inhibition of peroxisomal catalase, concomitantly with an increase of ROS, is known to result in mitochondrial redox imbalance^38^. Moreover, Env is known to induce the mitochondrial pathway of apoptosis in bystander CD4+T cells^39^. In addition, a recent study showed that a subset of miRNAs, targeting peroxisomal proteins, are up-regulated in the brain of HIV/AIDS patients with and without HAND (HIV-associated neurocognitive disorders). The authors found that these miRNAs significantly decreased peroxisome number and affect their morphology^40^. In this context, we focused our work on peroxisomes and demonstrate that these organelles are targeted by Env-induced autophagy. Our results suggest that peroxisomes are insufficiently recognized targets of HIV-1 infection and open new routes in the characterization of the relationships between autophagy and this virus. Obviously, we cannot exclude a role of the mitochondria and the endoplasmic reticulum in our model and it would be important to test if these organelles, as well as other antioxidant pathways, are also targeted by autophagy in the same context.

By analyzing both peroxisomal proteins and entire peroxisomes, we were able to demonstrate, for the first time, that pexophagy plays a major role in a viral infection, probably by impairing the control of induced oxidative stress. Pexophagy, which occurs at the cellular basal level, is thus induced upon Env exposure. It would be important now to decipher the specific Env-induced signalling pathway triggered upstream of this phenomenon and particularly the role of the kinase ATM (ATM serine/threonine kinase). Indeed, this protein is activated by peroxisomal ROS, leading to pexophagy and involves the selective targeting of PEX5 by p62/SQSTM1^41^.

Concerning HIV-1 infection, the degradation of cellular antioxidant system requires a special attention as a known side-effect of ART (Anti-Retroviral Treatment) is to induce an oxidative stress in patients^42–45^. Moreover, oxidative stress is also clearly involved in the development of HIV-1-associated disorders such as neurotoxicity, dementia and immune imbalance^46–51^. Several HIV-1-associated co-morbidities related to an increased risk of cardiovascular diseases development are favoured by the oxidative stress induced by the presence of the virus, such as high blood pressure, atherosclerosis and myocarditis^52–54^. Moreover, in this infectious context, the peroxisome depletion could be beneficial for the pathogens and its escape from innate immunity. Indeed, peroxisomes have been identified as antiviral innate immunity signalling platforms since they harbour MAVS (Mitochondrial AntiViral Signaling proteins) at their surface and participate to type I interferon production in response to viral infection^55^. Beyond this, one of the major barriers against the eradication of HIV-1 in patients is the presence of cellular reservoirs that contain integrated, replication-competent provirus, resistant to ART. Even if these treatments maintain a very low level of viral replication and have considerably ameliorated the HIV-1 infected patient life, a rebound of viremia is observed upon treatment interruption. Interestingly, a new therapy, based on the oxidative stress response, has been proposed to target the HIV-1 reservoir. The AuranoFin (AF) is a gold compound with a pro-oxidant function^56^ capable of inducing apoptosis of memory T cells based on their oxidative status^57^. AF combined with ART has been shown to completely suppress SIV (Simian Immunodeficiency Virus) viremia in macaques. More interestingly, the survival of treated macaques is maintained for 2 years after the treatment interruption without any sign of immunodeficiency^58^. These results indicate that apoptosis-inducing drugs, acting on oxidative stress, could be envisaged to reduce the HIV-1 reservoir. It would be thus interesting to analyze if AF, combined to ART, could modulate memory T cells autophagy.

In conclusion, our results open new route for future investigations at multiple levels of HIV-1-host interactions and, to our knowledge, they demonstrate for the first time a role of pexophagy during a viral infection.

## Materials and methods

### Cell culture

The HEK293, HEK.Env^39^ and HEK/CD4.403/CXCR4^13^ cell lines were cultured in DMEM supplemented with 1% penicillin/streptomycin, 1% Glutamax (Life technologies), and 10% Fetal Calf Serum (FCS, Sigma-Aldrich). The CEM T cell line was provided by the ATCC. The CEM-derived A2.01/CD4.403 cell clone, which expresses a mutant form of CD4 truncated at position 403, was already used in^13^ and was provided by D.R. Littman (New York Medical College, New York, New York, USA). The 8.E5 cell line, a CEM-derived T cell line containing a single integrated copy of HIV-1 and unable to produce infectious virions, was provided by F. Barré-Sinoussi (Institut Pasteur, Paris, France). These T cell lines were cultured in RPMI 1640 medium supplemented with 1% penicillin/streptomycin, 1% Glutamax, and 10% FCS. Primary CD4+ T cells were purified from the blood of healthy donors by negative selection using the Human CD4+ T Cell Enrichment cocktail (StemCell Technologies). These primary cells were cultured in complete RPMI 1640 medium.

### Reagents and antibodies

AMD3100, NAC, Spautin 1, E64d, Pepstatin A, Bafilomycin A1 were purchased from Sigma-Aldrich. The anti-PEX14 and anti-BECLIN 1 antibodies (Abs) were purchased from Proteintech (references 105941-AP and 11306-1-AP, respectively). The anti-p62/SQSTM1 Ab was purchased from Euromedex (reference GTX100685). The anti-catalase Ab was purchased from Abcam (references ab1877-10). The anti-LC3B and the anti-GAPDH Abs were purchased from SIGMA-ALDRICH (reference L7543 and G9295, respectively).

### Transfections

*RNA interference:* ON-TARGET plus Human SMART Pool from Dharmacon were used for RNA interference targeting *Beclin 1* (reference L-010552) and siGENOME Human SQSTM1 siRNA – SMART Pool for *p62/SQSTM1* (reference M-010230). The ON-TARGET plus Non-targeting Pool (reference D-001810-10) was used as a control. Briefly, 0.5×10^6^ target cells were transfected using LipoRNAiMax (Invitrogen, P/N56531) with 20nM siRNA according to the manufacturer’s instructions. 16h after transfection, target cells were co-cultured with effector cells expressing, or not, Env (1.10^6^/ml) for 48h before analysis.

*Plasmids:* 0.5×10^6^ target cells were cultured in 6-well plates and transfected with pEGFP-SKL (Addgene#53450 from Jay Brenman Lab) plasmids using TurboFect Transfection Reagent (Thermo-Scientifics R0531) according to the manufacturer’s instructions. Transfected target cells were co-cultured with effector cells expressing or not Env for 48h before analysis.

### RT-PCR

Target cells were co-cultured with effector cells expressing, or not, Env for 48h. RNA was purified from 10^6^ target cells using the Nucleospin RNA plus (Macherey-Nagel), and RT-PCR was performed using the QIAGEN OneStep RT-PCR kit according to the manufacturer’s instructions. The oligonucleotides used *Pex14* and *GAPDH* mRNA are: *Pex14* FOR: 5’-GGGCTGACAGATGAAGAGATTG-3’ and *Pex14* BACK: 5’-CCTGGATCTTCTGCTGCTGCTG −3’ GAPDH FOR: 5’-CCCATCACCATCTTCCAG-3’; GAPDH BACK: 5’-CCTGCTTCACCACCTTCT-3’.

### Apoptosis analysis

Target cells were co-cultured with effector cells expressing, or not, Env for 48h. Apoptosis was analyzed in the target cells using “Cleaved PARP (Asp214) Cellular Kit” from CisBio according to the manufacturer’s instructions. FRET analysis was measured using a SPARK 10M TECAN.

### ROS analysis

Target cells were co-cultured with effector cells during 48h. Then, cells were incubated for 15min at 37°C in HBSS containing 0.86 mM DHR 123 (ThermoFisher, reference D632) and fluorescence intensity was rapidly measured at 543 nm by flow cytometry.

### Western blot analysis

Target cells were co-cultured with effector cells during 48h. Target cells were then harvested and lysed directly in Laemmli buffer 2x and heated for 10 minutes at 95°C. Cell lysates were loaded on 12% Prosieve 50 gels (Lonza) and transferred to polyvinylidene fluoride (PVDF) membranes. After a blocking step in PBS containing 5% casein for 1h at room temperature, blots were incubated overnight at 4°C with the primary Ab in the blocking buffer. After 3 washes with PBS+0.05% Tween, the blots were incubated for 1h at room temperature with peroxidase-coupled antiserum diluted in blocking buffer. After further washes, the immune complexes were revealed by ECL (Clarity Western ECL Substrate from BioRad, reference 170-5061) and analyzed with a ChemiDoc camera (Bio-Rad). Quantification of protein expression was performed using the Image lab Software (Bio-Rad). Data were normalized with reference to the densitometry analysis of GAPDH.

### Immunofluorescence studies

Target cells were cultured on glass slides and co-cultured with effector cells during 48h. For the staining, target cells were first incubated with 1% PFA for 10min and then, with methanol during 5min. After a saturation step in a PBS/0.1%Saponin/1%FBS buffer, the primary Ab was incubated during 2h at room temperature. After several washes, the secondary Ab coupled with a fluorochrome was added during 1h. Colocalization analysis was done using a Leica SP5 confocal microscope. Acquired images were analyzed using the Fiji plugin JaCOP. The Mander’s coefficient, measuring the overlap of GFP-SKL in LC3B staining, was determined on each z-stack of each acquisition before being averaged on all stacks of each cell^59^. For the « Cell profiler » analysis, a procedure was set up to discriminate the peroxisome over their area size.

### DAB Staining

Target CD4+ T cells were harvested after 48h of co-culture with effector cells and, then, incubated in RPMI medium for 3h at 37°C on poly-lysine coated glass slides. After several PBS washes, cells were fixed during 10min in 4% PFA-containing 0.2% Triton. Slides were then incubated during 15min in a DAB containing buffer (Sigma-Aldrich, reference D4293) according to the manufacturer’s instructions. After 3 washes with water, the DAB precipitates were visualised by light microscopy.

### Statistics

Differences were considered significant at *p < 0.05, **p < 0.01, and ***p < 0.001. Variances were measured using the Wilcoxon test on GraphPad Prism.

## Abbreviations

ROS: Reactive Oxygen Species
HIV-1: Type 1 Human Immunodeficiency Virus
Env: HIV-1 envelope glycoproteins
BafA1: Bafilomycin A1
NAC: N-Acetyl-Cystein
AP: AntiProteases
PTS: Peroxisomal Targeting Signal
DAB: 3’3-DiAminoBenzidine

## Supplementary Figures

**Supplementary Figure S1:** Catalase expression level was analyzed in target CD4+ T cells after 48h of co-culture with effector HEK.293 cells expressing or not Env, in the presence or absence of AMD3100 (1μg/ml). The expression level ratio of catalase was calculated between the conditions with or without Env and normalized to that obtained with anti-GAPDH Ab.

**Supplementary Figure S2:** Catalase expression level was analyzed in target CD4+ T cells after 48h of co-culture with effector HEK.293 cells expressing or not Env, in the presence or absence of AP (E64d + Pepstatin A, 10 μM each). The expression level ratios of catalase were calculated between the conditions with or without Env and normalized to that obtained with anti-GAPDH Ab.

**Supplementary Figure S3:** Pex14 mRNA level was analyzed by RT-PCR in target CD4+ T cells after 48h of co-culture with effector HEK293 cells expressing or not Env, in the presence or absence of AP (E64d + Pepstatin A, 10 μM each). GAPDH mRNA was used as a control.

**Supplementary Figure S4:** HEK/CD4.403/CXCR4 cells were co-cultured during 48h with effector cells expressing or not Env (8.E5 or CEM, respectively) after their transfection with *Beclin 1* (siBEC1) or *control* (siCT) siRNAs. The expression level ratios of catalase was calculated between the conditions with or without Env and normalized to that obtained with anti-GAPDH Ab.

**Supplementary Figure S5:** Target CD4+ T cells were harvested after 48h of co-culture with effector cells in the presence or not of AP (E64d + Pepstatin A, 10 μM each). The cells were then fixed during 10min in 4% PFA-containing 0.2% Triton. Cells were incubated during 15min in a DAB containing buffer and precipitates, due to peroxidase activity, were visualised by light microscopy.

