## Supplementary figures and images for "HIV-1 Env induces pexophagy and an oxidative stress leading to uninfected CD4+ T cell death"

### Supplemental Figures

Supplemental Figures

S1

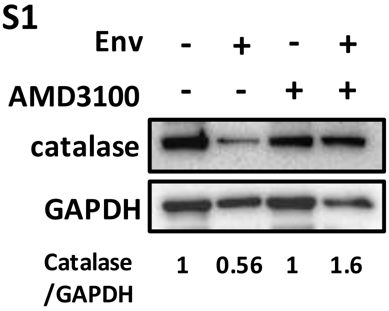


S2

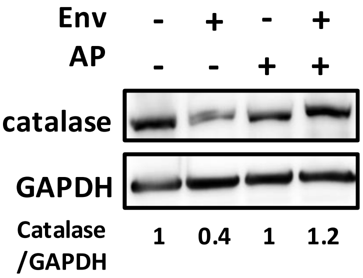


S3

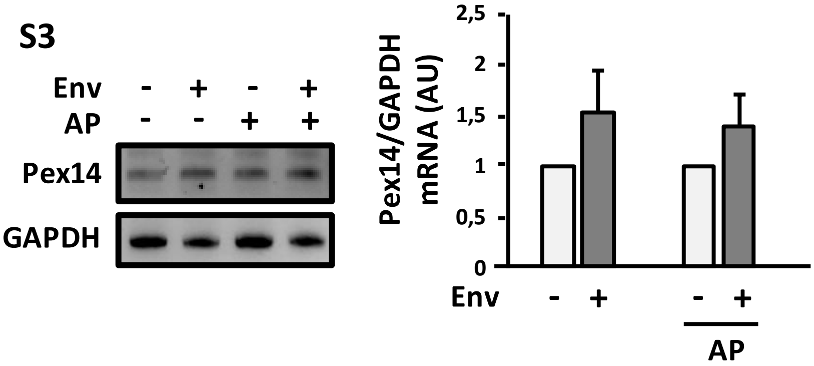


S4

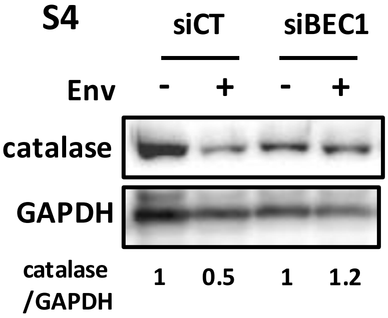


S5

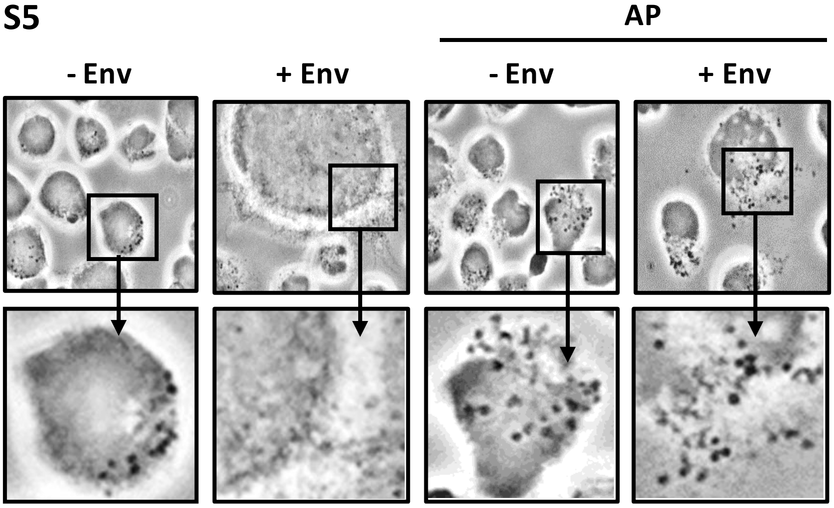
